# DSCN: double-target selection guided by CRISPR screening and network

**DOI:** 10.1101/2021.09.06.459081

**Authors:** Enze Liu, Xue Wu, Lei Wang, Yang Huo, Huanmei Wu, Lang Li, Lijun Cheng

## Abstract

Cancer is a complex disease with usually multiple disease mechanisms. Target combination is a better strategy than a single target in developing cancer therapies. However, target combinations are generally more difficult to be predicted. *C*urrent CRISPR-cas9 technology enables genome-wide screening for potential targets, but only a handful of genes have been screend as target combinations. Thus, an effective computational approach for selecting candidate target combinations is highly desirable. Selected target combinations also need to be translational between cell lines and cancer patients.

We have therefore developed **DSCN (<u>d</u>ouble-target <u>s</u>election guided by <u>C</u>RISPR screening and <u>n</u>etwork)**, a method that matches expression levels in patients and gene essentialities in cell lines through spectral-clustered protein-protein interaction (PPI) network. In DSCN, a sub-sampling approach is developed to model first-target knockdown and its impact on the PPI network, and it also facilitates the selection of a second target. Our analysis first demonstrated high correlation of the DSCN sub-sampling-based gene knockdown model and its predicted differential gene expressions using observed gene expression in 22 pancreatic cell lines before and after MAP2K1 and MAP2K2 inhibition (*R*^2^ = 0.75). In our DSCN algorithm, various scoring schemes were evaluated. The ‘diffusion-path’ method showed the most significant statistical power of differentialting known synthetic lethal (SL) versus non-SL gene pairs (*P* = 0.001) in pancreatic cancer. The superior performance of DSCN over existing network-based algorithms, such as OptiCon[1] and VIPER[2], in the selection of target combinations is attributable to its ability to calculate combinations for any gene pairs, whereas other approaches focus on the combinations among optimized regulators in the network. DSCN’s computational speed is also at least ten times faster than that of other methods. Finally, in applying DSCN to predict target combinations and drug combinations for individual samples (DSCNi), we showed high correlation of DSCNi predicted target combinations with synergistic drug combinations (*P* = 1e-5) in pancreatic cell lines. In summary, DSCN is a highly effective computational method for the selection of target combinations.

**Author Summary:** Cancer therapies require targets to function. Compared to single target, target combination is a better strategy for developing cancer therapies. However, predicting target combination is much complicated than predicting single target. Current CRISPR technology enables whole genome screening of potential targets. But most of the experiments have been conducted on single target (gene) level. To facilitate the prediction of target combinations, we developed DSCN **(<u>d</u>ouble-target <u>s</u>election guided by <u>C</u>RISPR screening and <u>n</u>etwork)** that utilize single target-level CRISPR screening data and expression profiles for predicting target combinations by connecting cell-line omics-data with tissue omics-data. DSCN showed great accuracy on different cancer types and superior performance compared to existing network-based prediction tools. We also introduced DSCNi derived from DSCN and designed specific for predicting target combinations for single-paitent. We showed synergistic target combinations predicted by DSCNi accurately reflected synergies on drug combination levels. Thus, DSCN and DSCNi have the potential be further applied in personalized medicine field.

## Introduction

The complexity of cancer is widely recognized, with heterogeneous disease mechanisms underlying primary, metastatic, and drug-resistant tumors [3, 4]. Therefore, translational cancer research now focuses on the identification of combinational rather than single targets and the selection of drug combinations instead of single drugs [5, 6]. Synthetic lethality (SL), a key concept in the simultaneous targeting of two genes that contribute to tumor vulnerability [7], requires the loss of both genes in a pair to be lethal to a cancer cell. A CRISPR-based double knockout (CDKO) system has recently been developed to effectively screen gene pairs or target combinations [8, 9]. In this paper, we will use the terms gene pair and target combination interchangeably because they represent equivalent concepts. Screening using the CDKO system, however, is limited by the number of genes to be screened. For instance, if we screen target combinations among 100 genes, and each gene has four gRNAs, there will be (4 × 100)^2^/2 = 80,000 combinations, a scale that is feasible in a CDKO system. However, across the genome, if we screen target combinations among 10,000 genes and select only one gRNA per gene, the resulting 10,000^2^/2 = 50,000,000 combinations will be practically infeasible. Therefore, a computational approach is very much needed to rank and select top candidate gene pairs for CDKO analysis.

The many computational methods developed to identify potential candidate SL gene pairs fall into two major categories: machine learning and statistical inference. The machine-learning approach has a much longer history, with the generation of large-scale double knockout data in yeast in 2004 [10]. Several methods, including multiple network decision tree [11], protein interaction network [12], and multi-network and multi-classifier [13] approaches, have demonstrated the significant predictive performance of SL gene pairs using features derived from network topology, gene ontology, and gene function sets. Recently, researchers applied a systems-biology framework called ‘mashup’ that allows for the acquisition and synthesis of data from diverse sources [14]. This method performed even better than the other network analyses in predicting SL gene pairs, further demonstrating the ability of such computational algorithms as random walk with restart to integrate and characterize the biological network topology successfully. Group-sparse collective matrix factorization (gCMF), another unique machine-learning method and recent major contribution to SL prediction [15], performed matrix factorizations among input data, such as gene expression, mutations, copy number variations (CNV), and CDKO, and identified a shared sub-matrix in which SL gene pairs can be predicted. Its performance was comparable to that of the mashup approach in several CDKO datasets derived from human cancer cell lines.

Statistical inference, on the other hand, relies on strong biological assumptions regarding the mechanisms of synthetic lethality. These methods infer SL gene pairs utilizing multi-omics data, such as CNV, mutations, gene expression, and single gene essentiality generated from CRISPR screening. In particular, CDKO data are NOT employed to train SL prediction in the statistical inference. DAISY is a notable early SL inference method [16], in which one primary assumption is that if the cancer cell is viable, the SL pair comprises one gene that is both active and essential if the other is inactive. MiSL (<u>mi</u>ning <u>s</u>ynthetic lethals), another statistical-inference approach [17], assumes that if one gene in an SL gene pair is inactive, the other must be active, and its activity is demonstrated through concordant changes in CNV and gene expression. Several other methods, such as ISLE (identification of clinically relevant synthetic lethality) [18], DiscoverSL [19], and ASTER (analysis of synthetic lethality by comparison with tissue-specific disease-free genomic and transcriptomic data) [20] were developed similarly, each making different biological assumptions regarding SL gene pairs.

Considering network topology, recent systems-biology-based statistical inference methods differed significantly from the other established SL statistical-inference methods. Here, we highlight two notable approaches, OptiCon (optimal control nodes) [1] and VIPER (virtual inference of protein activity by enriched regulon analysis) [2]. Both approaches primarily utilize gene-expression data to construct a biological network and rank and select target combinations that demonstrate optimal control of the network. OptiCon relies on a protein-protein interaction (PPI) network and models both signaling transduction and gene regulation during the selection of target combinations, and VIPER focuses on a gene-regulatory network model derived from mutual information among genes. Both approaches assume that the more a combination of two gene targets controls the network, the more likely those targets will be an SL pair.

Both statistical inference and machine learning have their advantages and disadvantages. A machine-learning approach optimizes prediction based on training data, but the quantity of training data limits the validity of its prediction of SL gene pairs. Extrapolation of the SL gene pairs from machine-learning prediction to other pathways will be challenging because most CDKO data generated from human cancer cells are sparse and biased toward several specific pathways. However, the prediction of genome-wide SL gene pairs using statistical-learning approaches relies strongly on biological assumptions and is technically unbiased. This can be particularly useful in the case of very limited CDKO data.

In this paper, we describe a new statistical inference method we have developed called DSCN (<u>d</u>ouble target <u>s</u>election guided by <u>C</u>RISPR screening and <u>n</u>etwork). It more resembles OptiCon and VIPER than other methods, such as DAISY or MiSL. Similar to OptiCon and VIPER, DSCN is built upon a biological network and transcriptome data, and the top combination targets are ranked and selected based on their control of or impact on the network.

DSCN differs from OptiCon and VIPER in its use of single-gene-library-based CRISPR-cas9 screening data, its focus on the overlapped networks between the cancer cell line and the corresponding primary tumor data, and most important, its consideration of the sequential selection of two targets, which involves the perturbation of transcriptome data for selection of the second target after selection of the first. This third model strategy, we believe, will make the selection of combination targets by DSCN more closely resemble the true biology. DSCN is also built upon our early research in the selection of single targets, SCNrank (<u>s</u>pectral <u>c</u>lustering for <u>n</u>etwork-based <u>ranki</u>ng) [21], which ranks and selects consensus single-gene targets between cancer cell lines and tumor samples.

## Materials and Methods

**Tables 1** details our data sources, including the types of cancer screened, data platforms and types, and sample numbers. We retrieved gene-expression and -mutation data for normal tissue and tumor samples for pancreatic and breast cancers from the Gene Expression Omnibus (GEO) [22, 23] and The Cancer Genome Atlas (TCGA) [24] and gene-expression and -essentiality data from the Cancer Cell Line Encyclopedia (CCLE) [13] and Project Achilles [25-27], downloaded PPI data from STRING [28], extracted drug-target data from DrugBank [12], and downloaded synthetic lethal gene-pair data from the SynlethDB database [29] and drug-sensitivity data from the DrugComb database [30].

**Table 1.** Datasets used in this study.

| **Part 1. Multi-omics data** |  |  |  |
| --- | --- | --- | --- |
| **Cancer type** | **Data platform** | **Data type** | **Data (n, sample size)** |
| Pancreatic cancer cell lines | Affymetrix U133 2.0 | Gene expression | GSE36133 (43), GSE46385 (7), GSE21654 (22), GSE17891 (20)<br>Total sample size = 92 |
|  | CRISPR screening | Gene essentiality | Project Achilles (v3.3.8)<br>Total sample size = 26 |
| Pancreatic tissue samples | Affymetrix U133 2.0 | Gene expression (tumor) | GSE42952 (33), GSE51978 (2), GSE16515 (36), GSE15471 (39), GSE23952 (3)<br>Total sample size = 113 |
|  | Affymetrix U133 2.0 | Gene expression (normal) | GSE46385 (3), GSE16515 (16), GSE15471 (39)<br>Total sample size = 58 |
|  | Illumina DNA-seq & RNA-seq | Mutation and gene expression (tumor) | TCGA ductal and lobular neoplasms (150), adenomas and adenocarcinomas (29) |
|  | Illumina RNA-seq | Gene expression (normal) | Solid tissue adjacent normal (41) |
| Breast cancer tissue samples | RNA-seq | Gene expression (tumor) | TCGA triple negative breast cancer sample (115) |
|  |  | Gene expression (normal) | TCGA triple negative breast cancer sample (163) |
| Breast cancer cell lines | Affymetrix U133 2.0 | Gene expression | GSE36133 (12) |
|  | CRISPR Screening | Gene essentiality | Project Achilles (v3.3.8)<br>Total sample size = 28 |
| **Part 2. Databases** |  |  |  |
| **Data type** | **Database** | **Data** |  |
| Protein-protein interaction (PPI) network | STRING[28] | PPI data in STRING database for human (v11): 11,609,230 interactions |  |
| Drug targets | DrugBank[31] | Food and Drug Administration (FDA)-approved drugs and their associated target proteins: 1,769 gene targets |  |
| Synthetic lethal pairs | SynlethDB[29] | 19,613 synthetic lethal gene pairs in human cancer |  |
| Drug sensitivity data | DrugComb[30] | Drug synergies among cell lines on 5,226 drug pairs (HS5 |  |

### Steps of DSCN algorithm

DSCN algorithm consists of six steps (**Figure 1**):

**Figure 1.**
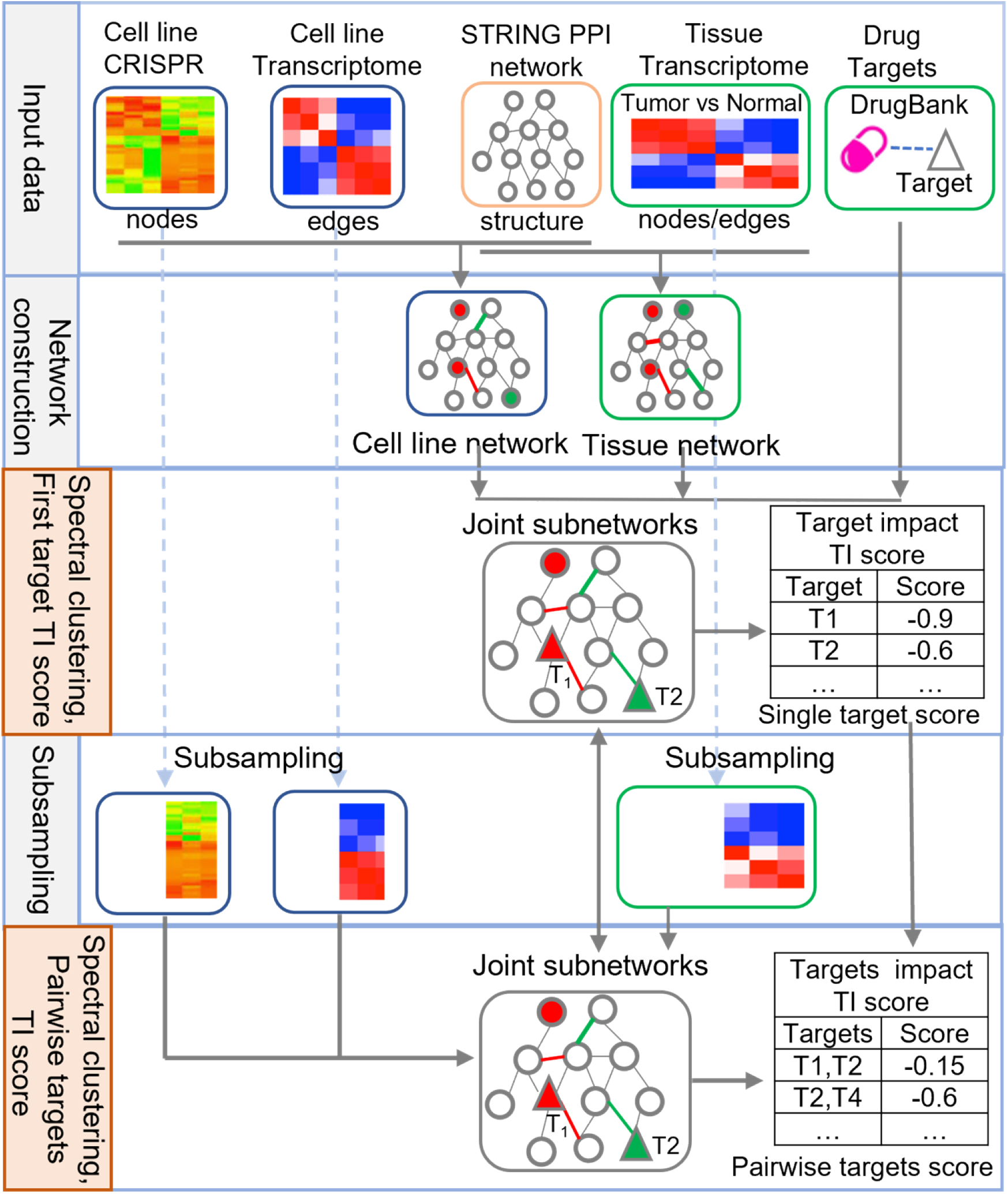
Overview of <u>d</u>ouble-target <u>s</u>election guided by <u>C</u>RISPR screening and <u>n</u>etwork (DSCN).

#### Step 1: Network construction

In this step, we construct two integrated function networks, a tissue network *G*_*t*_ and a cell-line network *G*_*c*_. *G*_*t*_ consists of a skeleton from the STRING PPI network and edge weights from gene pair-wise Pearson correlations in tumor samples, and node weights are the fold changes in gene expression between tumors and normal tissue. A high fold change indicates higher gene expression in the tumor than in normal tissue. Assume that there are a total of n genes (nodes) in *G*_*t*_. The affinity matrix *S*_*t*_ denotes the edge weights, and diagonal matrix *D*_*t*_ denotes the node weights in Equation (1):

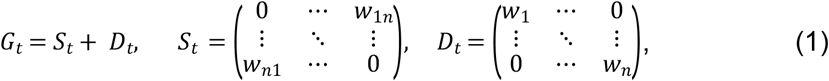

where *w*_ab_, *a* ≠ *b* ∈ (1,*n*) in *S*_*t*_ indicates the edge weight (correlation) between genes a and b in the tissue network; and *w*_*i*_ in *D*_*t*_ is the tumor versus normal fold change in the expression of gene *i, i* = 1,…,*n*.

Similarly, *G*_*c*_ consists of an identical skeleton from the same STRING PPI network and edge weights from pair-wise gene correlations in cell-line samples. Unlike *G*_*t*_, the node weight of *G*_*c*_ is from CRISPR-Cas9 screening data, which is indicated as the gene essentiality value. The gene essentiality value can be generally interpreted as the fold change in cell count before and after gene knockout. Genes demonstrating smaller fold change are more essential. In this study, all the essentiality values are log2 transformed. Similarly, *G*_*c*_ is decomposed into affinity matrix *S*_*c*_ for edge weight and diagonal matrix *D*_*c*_ for node weight in the cell-line network *G*_*c*_ = *S*_*c*_ + *D*_*c*_.

#### Step 2: Construction of Laplacian matrices for the tissue and cell-line networks

A Laplacian matrix measures all properties of a network, including node weight, edge weight, and connectivity. In this second step, we construct Laplacian matrices for the tissue network *G*_*t*_ and cell-line network *G*_*c*_ as:

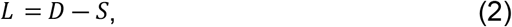

in which *D* is the diagonal matrix and *S*, the affinity matrix, defined in Equation (1), and *L*_*t*_ is the Laplacian matrix for the tissue network and *L*_*c*_, that for the cell-line network.

#### Step 3: Spectral clustering for tissue network

We perform spectral clustering only on the Laplacian matrix of the tissue network *L*_*t*_ as:

I. Normalize the Laplacian matrix *L*_*t*_ to *L*′_*t*_:

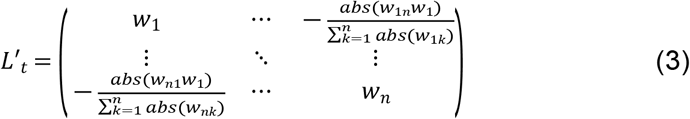

In the normalized Laplacian matrix *L*′_*t*_, all diagonal elements are positive, and all other elements are negative. The row sum of non-diagonal elements is equal to its corresponding diagonal.
II. Perform eigen decomposition for matrix *L*′ to obtain the spectrum *E* = {*λ*_1_, *λ*_2_…*λ*_*n*_}, where 0 = *λ*_1_ ≤ *λ*_2_ ≤ … ≤ *λ*_*n*_, and their corresponding eigenvector.
III. Choose the *k* smallest non-negative eigenvalues {*λ*_*i*_,…,*λ*_*i*+*k*_} and their corresponding eigenvectors, and combine these *k* eigenvectors into an *n* × *k* matrix, *H*.
IV. In this *H* eigenvector matrix, each row represents a gene node, and *k* columns represent the coordinate values of a gene node. The row vectors in *H* are used to calculate the Euclidean distance between a pair of gene nodes. We then perform K-means clustering for *n* nodes. To select the number of clusters, K’, to produce a good fit, we calculate Hartigan’s number, which measures the quality of clustering results. We select the optimal K’ and constrain it further to less than 10 for practical consideration. This spectral clustering leads to K’ exclusive clusters (i.e., subnetworks). From the tissue network *G*_*t*_, subnetworks 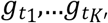 are classified.

#### Step 4: Mapping the tissue/cell-line network and calculating the impact score of Target 1

The cell-line network *G*_*c*_ is then mapped to the spectral clusters, 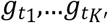, generated from tissue network *G*_*t*_ in Step 3. Because tissue network *G*_*t*_ and cell-line network *G*_*c*_ share the identical network structure, i.e., nodes and connections, *G*_*t*_ subnetworks, 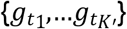 are mapped to *G*_*c*_ subnetworks 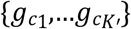 using their common node names and connections.

The target impact score will be calculated based on the cell-line subnetworks 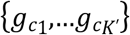 We focus on all Food and Drug Administration (FDA)-approved drug targets (see **Table 1**) to calculate our target score. The impact score of a target 1 (*T*1) is calculated as the sum of the impact score itself and its impact on the rest of the genes in the network. Its general form is defined in Equation (4):

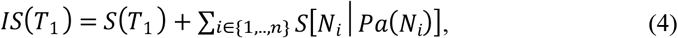

in which, {*N*_*i*_, *i* = 1,…*n*} are the gene nodes in the network other than *T*1, and *Pa*(*N*_*i*_) is a set of parent nodes of *N*_*i*_. In particular, the impact score on *N*_*i*_ depends on its parent nodes, *Pa* (*N*_*i*_). **Figure 2** illustrates the three different methods of calculating the impact score–the most-probable, random-walk, and diffusion paths.

**Figure 2.**
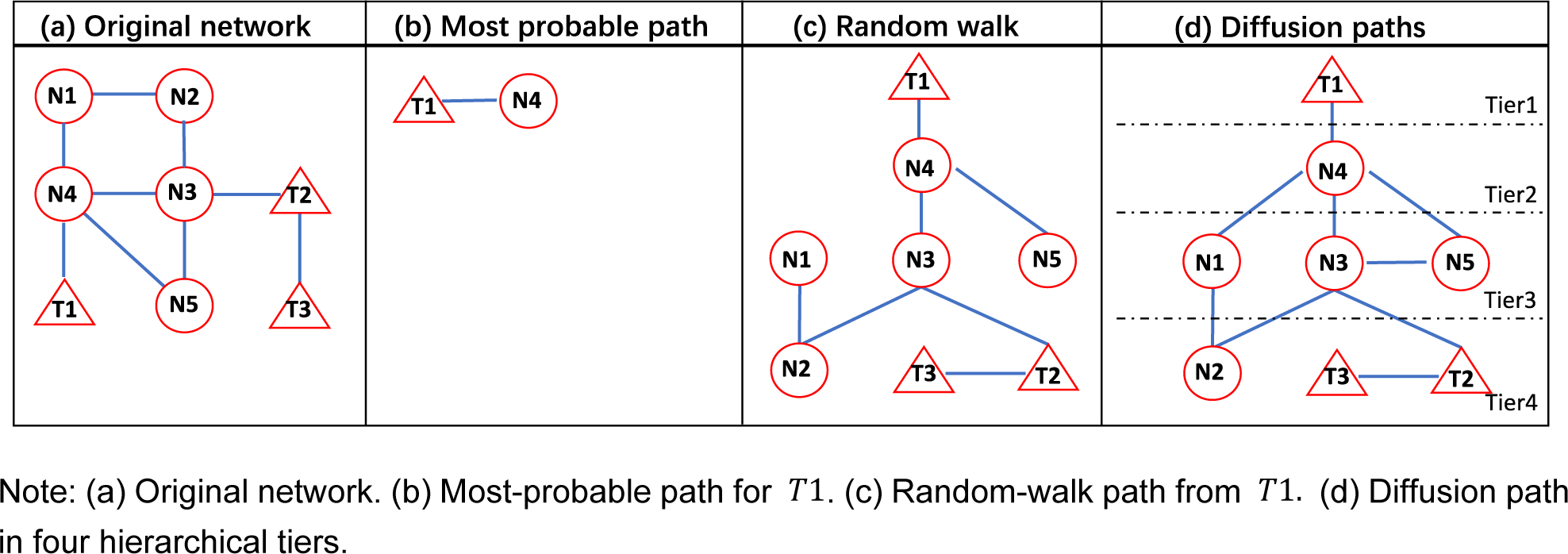
Network configurations for three methods to calculate impact score.

##### Most-probable path

The immediate children of *T*1 are the gene nodes directly connected to *T*1, e.g., N4 is the direct child *T*1 in **Figure 2b**. In this method, we will count only the immediate children of *T*1 in calculating the impact score. Without loss of generality, let *ch*(*T*1) be the set of immediate children of *T*1. The most probable path of *T*1 is the one that has the smallest impact score among *ch*(*T*1). Based on the general impact score as calculated in Equation (4), the most-probable-path impact score is defined in Equation (5):

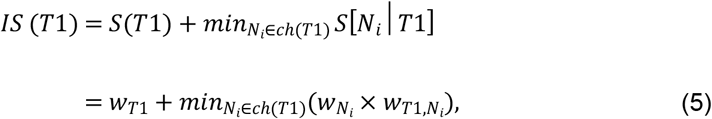

where *w*_*T*1_ and 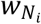 indicate their node weights, and 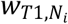 indicates their edge weight.

##### Random walk path

The random-walk score is calculated in two steps. Step 1 is a random walk in the network, in which the random walk has a transition probability of traveling from one node to another. In **Figure 2c**, starting from *T*1, each node *N*_*i*_ is randomly visited. Here we used normalized edge weight for transition probability as defined in Equation (6):

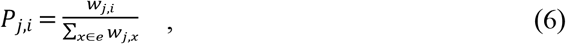

where *P*_*j,i*_ is the transition probability from *N*_*j*_ to *N*_*i*_, *w*_*j,i*_ is the edge weight between them, and ∑_*x*∈*e*_ *w*_*j,x*_ is the sum of all edge weights of *N*_*j*_. In this Markov process, a node can be visited multiple times. We set the total number of random-walk steps as 2*n*, where *n* is the total number of nodes in the network.

Then, in Step 2, we defined the parent node as the node that visited *N*_*i*_ first, i.e., *Pa*(*N*_*i*_). Hence, the impact score of *T*1 becomes:

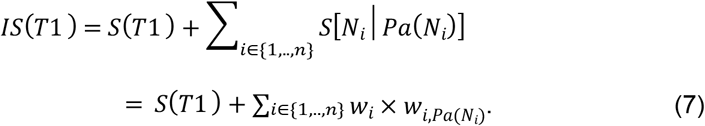

##### Diffusion path

Starting from *T*1, each node is visited in a hierarchical order. Therefore, the parent nodes of a node, *N*_*i*_, can be from the upper tier, i.e., *UpperTier* (*N*_*i*_), or the same tier, i.e., *SameTier* (*N*_*i*_). For instance, in **Figure 2d**, there are four tiers in the hierarchical structure starting from *T*1. The impact of *T*1 transmits from Tier 1 to Tier 4 in the network. Therefore, the impact score is defined in Equation (8):

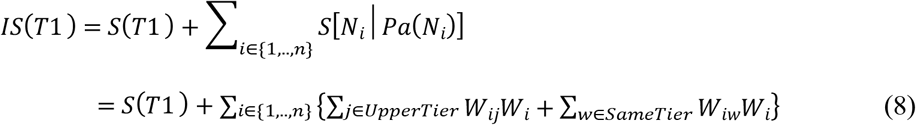

#### Step 5: Subsampling and Target 2 (*T2*) score and selection

Once *T1* is selected, we remove cancer cell lines with higher expression of the *T1* than its sample mean and only keep cell lines with its expression lower than mean.This subsampling method characterizes the knockdown of the *T1*. Similarly, we also remove cancer cell lines with higher *T1* essentiality scores than the sample in our subsampling. After the resampling, we construct the cell-line network *G*_*c*_ as Equation (2) using the subsampled cell-line subsamples. We follow the same **Step 3** in mapping *G*_*c*_ to 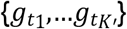 and calculate the *T2* impact score following the same algorithms defined in **Step 4**. The *T2* impact score is then denoted as *IS* (*T*2|*T*1), because the subsampling and network depend on *T*1.

#### Step 6: Calculation of impact score for target combinations

Because *T*1 and *T*2 and their impact scores are computed sequentially, the combinational impact score will consider both sequential orders in Equation (9), in which *T*1 ≠ *T*2:

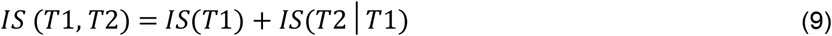

#### Tissue cell-line subnetwork similarity measure

We measure the similarity of each subnetwork pair 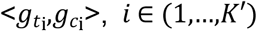 between tissue and cell-line using the following scheme:

##### I. Normalization of node weight (diagonal)

To make two subnetworks, 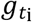 and 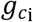, comparable, we normalize the cell-line diagonal matrix 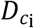 according to the tissue diagonal matrix 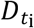 using the following formula:

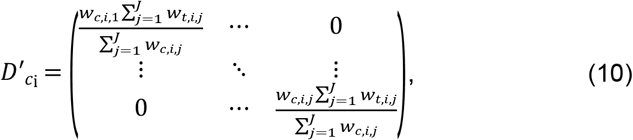

in which *w*_*c,i,j*_ denotes the node weight *j* ∈ (1,*J*) in the cell-line subnetwork, and *w*_*t,i,j*_, that in the tissue subnetwork. *J* is the total number of nodes in 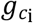 and 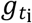.

##### II. Normalization of edge weight

The Laplacian matrices for each subnetwork pair, 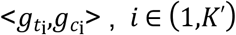, are defined similarly as Equation (3): 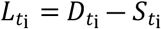 and 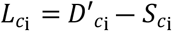. After node-weight normalization, trace 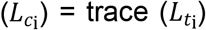. Then, their edge weights (non-diagonal elements) are normalized accordingly using the formula:

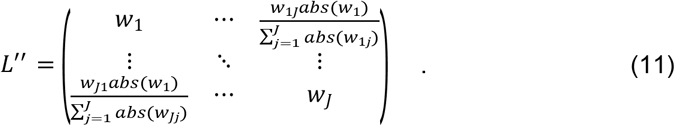

Until this step, all edges (non-diagonal elements) in both Laplacian matrices, 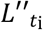 and 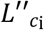, acquired node features during normalization. We keep the original directions (positive or negative) of node weights and edge weights for the following distance calculation.

##### III. Distance calculation

For two corresponding subnetworks 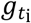 *and* 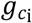 in tissue and cell-line, we calculate the distance using their normalized Laplacian matrices 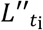 and 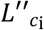:

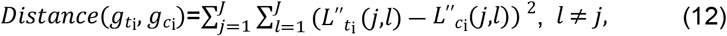

where L′′ (*i,j*) *i* ≠ *j* indicates the edge weight between nodes *l* and *j* in a given Laplacian matrix, and 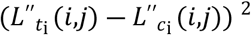 indicate the Euclidean distance between the same edges in two Laplacian matrices.

#### Construction of a DSCN algorithm for an individual cancer cell-line sample (DSCNi)

We apply DSCNi algorithm for scoring target combinations in a single cancer cellline for a single patient. Very similar to DSCN, in building up *G*_*c*_, DSCNi relies on a set of expression profiles for a cancer cell line to calculate the edge weights (i.e., correlations) between gene nodes. However, unlike DSCN, DSCNi uses a cell-line-specific essentiality score for node weights. Its impact score calculation for *T1, IS*(*T*1), follows exactly from **Steps 1, 2, 3, and 4**. In modeling the knockdown of *T1* in the subsampling in **Step 5**, we maintain the same *T1* subsampling as DSCN, i.e., we remove samples with higher expression of T1 than its sample mean. However, we will keep the same essentiality score for this individual cancer cell-line sample to calculate the Target 2 impact score. We calculate the final combination target impact score similarly as in DSCN, such that it has a comparable meaning to that calculated from DSCN.

### Analysis of association between drug- and target-combination synergy

The Bliss score [32] measures the synergistic effect of a drug combination, i.e., the effect of the drug combination on cell viability rather than the additive effects of its two component drugs. A two-drug combination is considered synergistic if its Bliss score exceeds 0.12 [33]. On the other hand, the target combination is predicted to be synergistic if the impact score of the drugs in combination is larger than the additive score of the constituent drugs, as in Equation (13), in which the impact scores of *IS*(*T*1, *T*2), *IS*(*T*1) and *IS*(*T*2) are calculated by (9) and (8):

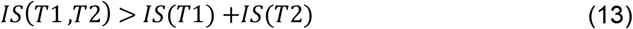

In this section, we will define an association analysis between drug-combination scores and target-combination synergy scores. Consider a cancer cell line screened by a set of drug combinations, and these drug combinations can be categorized as either synergistic or non-synergistic based on their Bliss scores. Then, for each drug combination, we identify all its two-target combinations, calculate their synergy scores, and classify the drug combinations as either synergistic or not as in Equation (13). In a 2 by 2 contingency table, the rows are drug synergy (Y/N), and columns are target synergy (Y/N). For each drug combination, all counts of target-combination synergy and non-synergy are added to the corresponding row with respect to drug-combination synergy or non-synergy. The association between drug- and target-combination synergy is tested using a Chi-square test.

## Results

### Validation of the subsampling scheme for determining the impact of target-gene knockdown in the DSCN algorithm

In the DSCN algorithm, we designed our subsampling method (**Step 5)** to model the impact of Target 1 knockdown in the cancer cell line. To demonstrate the validity of this sampling scheme, we identified a GEO dataset, GSE45757, that provided transcriptome profiles across 22 pancreatic cell lines before and after MAP2K1 and MAP2K2 inhibition. Our analysis focused on 1,301 neighbor genes of MAP2K1 and MAP2K2 in the PPI network. Using the subsampling approach, we calculated the log-fold changes in these 1,301 genes between groups with either high or low expression of MAP2K1 and MAP2K2 group, which represent the predicted impact of Target 1 knockdown in the subsampling scheme. On the other hand, the observed log-fold changes in these 1,301 gene expressions were calculated during MAP2K1 and MAP2K2 inhibition. **Figure 3** shows a strong correlation, *R*^2^ = 0.75, between the predicted and observed fold changes among these 1,301 neighbor genes of MAP2K1 and MAP2K2. Findings of this analysis strongly support subsampling as a valid model for determining the impact of target-gene knockdown.

**Figure 3.**
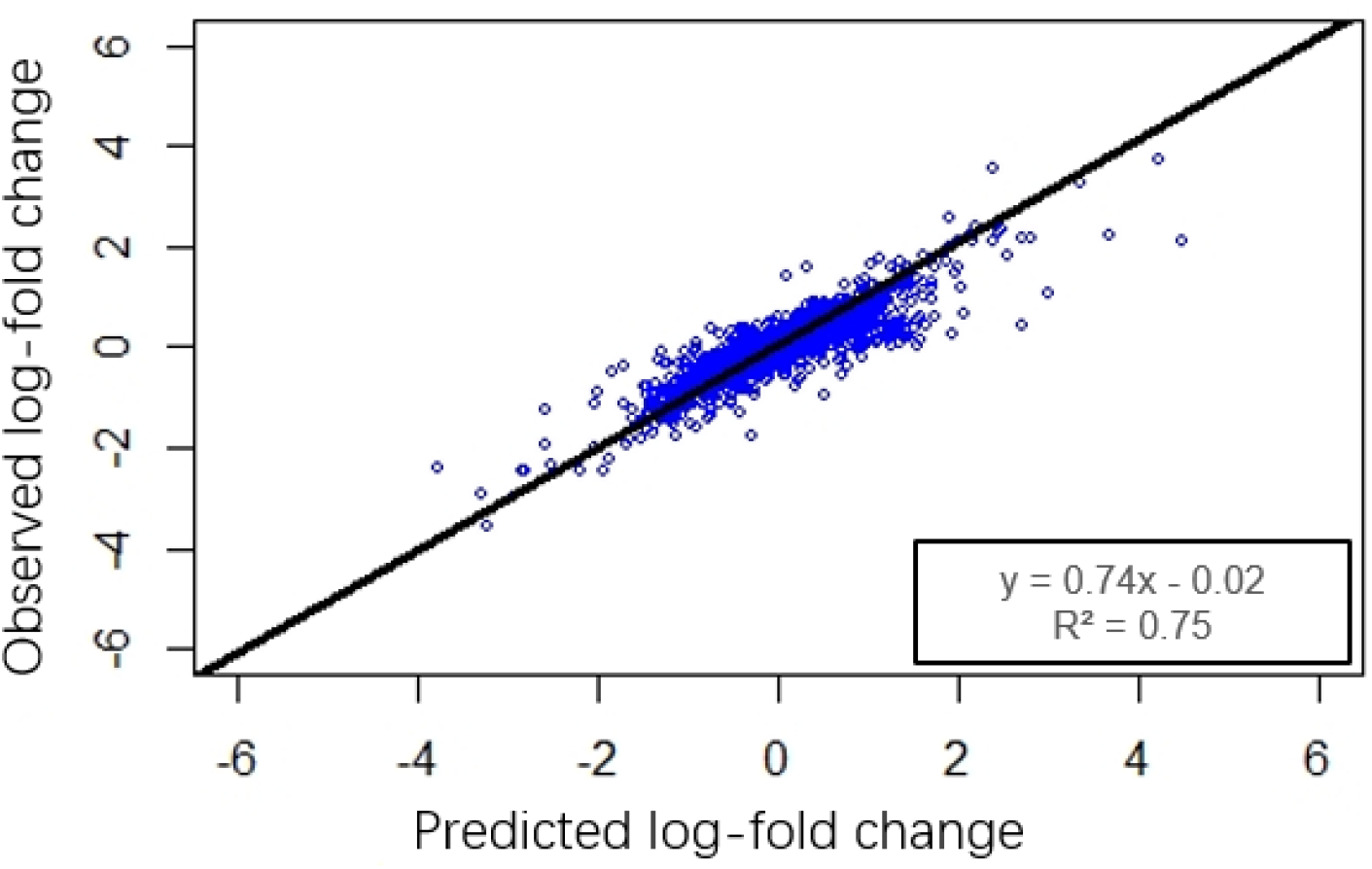
Correlation between the predicted and observed log-fold changes in gene expression among MAP2K1 and MAP2K2 neighbor genes in the protein-protein interaction (PPI) network.

### Comparison of impact scores of target combinations using known synthetic lethal gene pairs in pancreatic cancers

We proposed three different scoring schemes to model the impact of target-gene knockdown on the network–those of the most-probable, random-walk, and diffusion paths. In addition, the impact score can be calculated based on either the global or local PPI network. The local PPI network is the product from spectral clustering of the whole genome PPI network (global network). To compare the performance of these impact scores, we used the 23 reported synthetic lethal pancreatic gene pairs in SynlethDB [29] as benchmarks. We compared impact scores between them and the other 164 gene pairs, which were derived from 21 unique genes among the 23 SL gene pairs. We constructed a tissue-function network using 153 tumor and 58 normal expression profiles of the pancreas from the GEO database (**Table 1**) and a cell-line function network using CRISPR screening data of 26 pancreatic cell lines from Project Achilles and 92 pancreatic tumor cell-line expression profiles from the GEO database (**Table 1**). All expression profiles are generated by Affymetrix U1332.0 microarray.

Smaller impact scores indicated the stronger impact of the gene knockdown on the network. Calculation of the impact scores using the local network generated from spectral clustering revealed significant difference in diffusion-path-based impact scores (IS) between synthetic and non-synthetic lethal gene pairs (*P*-values) as well as lower impact scores of synthetic than non-synthetic lethal gene pairs. We observed the same trends with the other two impact scoring schemes, the most-probable and random-walk paths, i.e., lower IS score in the synthetic than non-synthetic lethal gene pairs that were not statistically significant.

Calculation of the impact scores using the global network and diffusion-path scoring scheme also yielded lower diffusion impact scores in the synthetic than non-synthetic gene pairs, though the differences were not statistically significant. The scores of the most-probable and random-walk paths, on the other hand, showed the reverse direction between synthetic and non-synthetic gene pairs. We therefore believe that using the diffusion-path and local networks, evaluation of the target-combination impact score is an ideal approach in selecting synthetic lethal gene pairs.

### Compare the selection of target combinations among DSCN, OptiCon, and VIPER

We compared the performance of DSCN with that of two existing algorithms for the selection of target combinations–OptiCon and VIPER. Both of these use transcriptome profiles to select combination targets, and their top target combinations are master regulators of synergy that have optimal control of their corresponding networks. OptiCon requires tumor transcriptome profiles and corresponding mutation data as input to infer master regulators and predict synergies among them, whereas VIPER uses transcriptome profiles from both tumor and normal samples to select regulons and infers synergies among the regulons. Because the pancreas microarray expression profile used in the previous section has no corresponding mutation information, we utilized pancreatic expression profiles in TCGA to construct a tissue function network. We used 179 pancreatic tumor expression profiles along with their mutation data and 41 adjacent normal expression profiles (Table 1). We also used expression profiles of 92 pancreatic tumor cell lines from GEO and CRISPR-screening data of 26 pancreatic cell lines from Project Achilles (Table 1). Together, these data served for benchmark comparison of the performance of the three algorithms.

In total, DSCN predicted 37,275 synergistic target combinations, OptiCon, 2,778, and VIPER,191. After mapping them onto all 12,821 synthetic lethal gene pairs within SynlethDB, neither OptiCon nor VIPER showed any overlap. However, DSCN demonstrated overlap of 936 target combinations. Among the 936 overlapped SL pairs, 79 were annotated as SL pairs specific to pancreatic ductal adenocarcinoma (PDAC). Of these 79, their predicted IS scores showed a0.34 Spearman correlation with their SynlethDB score (*P* < 0.01), and the predicted IS scores were significantly lower than that of 6,162 random combinations on t-test (*P* = 0.05). These 6,162 random combinations were derived from genes in the 79 SL pairs, but 79 were removed.

These benchmark comparison analyses were performed on Indiana University’s supercomputer, ‘Carbonate’. DSCN completed its search of target combinations on the single central processing unit core in 12 hours, a significantly faster speed than those using OptiCon (320 hours) and VIPER (141 hours). OptiCon mainly performs two computational tasks, calculating subnetworks and null distributions, each of which uses about 160 hours. Most ofthe computational time of VIPER, on the other hand, involves the generation of the transcriptome mutual information network using ARACNe [34], a classical tool of reconstructing regulatory network by calculating pair-wise mutual information between genes.

### Top-ranked target combinations and their associations with overall survival in patients with pancreatic cancer

We used expression profiles of tissues and cell lines from the GEO database (**Table 1**) to construct function networks and predict impact scores. Our dataset consisted of expression profiles of 153 tumors and 58 normal pancreas samples from GEO, CRISPR-screening data of 26 pancreatic cell lines from Project Achilles, and 92 pancreatic tumor cell-line expression profiles from the GEO database. This yielded 14,066 overlapped genes.

In this analysis, we focused on 1,437 drug targets of all FDA approved drugs in DrugBank and calculated their possible target combinations. Most interestingly, all genes in the top 230 target combinations are within the same subnetwork–the PDAC tissue subnetwork (**Supplementary Figure 1 A**) and cell-line subnetwork (**Supplementary Figure 1 B**). **Supplementary File 1** includes the full list of genes in this subnetwork.

**Table 2** displays the nine top-ranked target combinations and their annotations. Their Kaplan-Meier curves (**Figure 5**) are generated using TCGA PDAC clinical annotations from the Gene Expression Profiling Interactive Analysis (GEPIA) database [35]. Patient samples are categorized into two groups based on a target combination in which both genes are expressed either above (i.e., high-2) or below their means (i.e., low-2). Using log-rank test and Cox proportional hazard model to analyze the association between the expression of a target combination (high-2 versus low-2) and overall survival of patients with PDAC, we observed significant survival difference (*P* < 0.05, Table 2) of three of the nine top-ranked target combination comparisons, (EGLN1, TRFC), (FRK, TRFC), and (XDH, TRFC), their overall survival was worse for patients with high expression of these two genes than those with low expression.

**Table 2.** Analysis of overall survival among the nine top-ranked target combinations in pancreatic ductal adenocarcinoma (PDAC).

| Gene 1 | Gene 2 | Impact score | Log rank *P*-value | Hazard Ratio (HR) | HR *P*-value | Pathways |
| --- | --- | --- | --- | --- | --- | --- |
| EGLN1 | TFRC | -255.12 | 0.02 | 2.00 | 0.02 | hypoxia, ferroptosis |
| MAP2K2 | TFRC | -255.05 | 0.08 | 1.60 | 0.08 | MAPK, ferroptosis |
| HPSE | TFRC | -255.01 | 0.19 | 1.50 | 0.20 | Metabolism, ferroptosis |
| PPIC | TFRC | -254.86 | 0.06 | 1.80 | 0.06 | Immune system, ferroptosis |
| FRK | TFRC | -254.86 | 0.04 | 1.80 | 0.05 | Immune system, ferroptosis |
| EGLN1 | COX7C | -254.79 | 0.84 | 1.10 | 0.85 | Hypoxia, metabolism |
| XDH | TFRC | -254.75 | 0.001 | 2.40 | 0.002 | Metabolism, ferroptosis |
| MAP2K2 | COX7C | -254.72 | 0.14 | 0.65 | 0.15 | MAPK, oxidative phosphorylation |
| FTL | TFRC | -254.71 | 0.10 | 1.60 | 0.10 | ferroptosis, ferroptosis |

**Figure 4.**
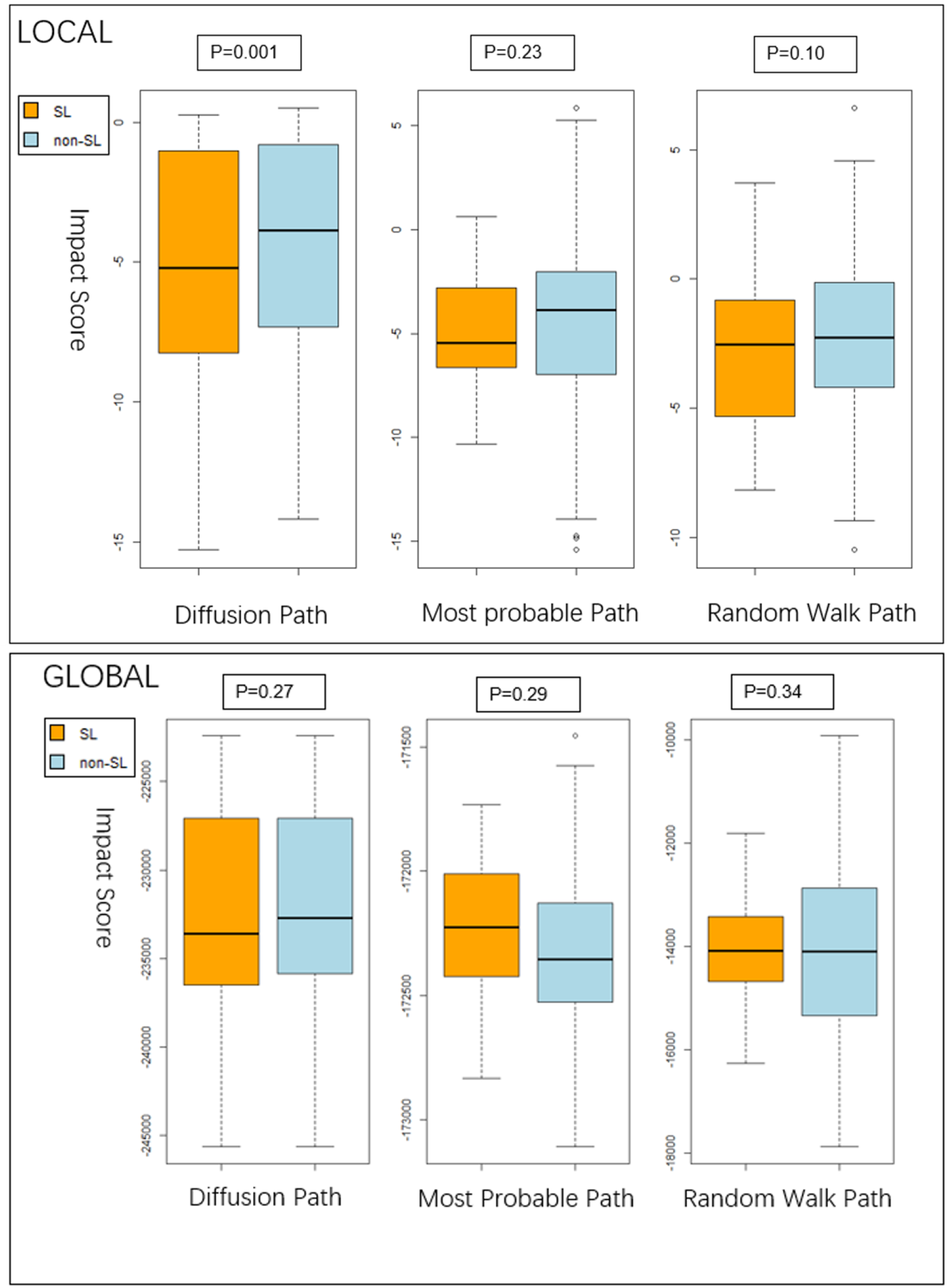
Comparison of target-combination impact scores using synthetic versus non-synthetic lethal gene pairs in pancreatic cancer. The three methods for calculating impact score–the most-probable, random-walk, and diffusion paths are defined in **Figure 2**. The impact scores (IS) are calculated from either the global protein-protein interaction (PPI) network (global) or the local PPI network (local).

**Figure 5.**
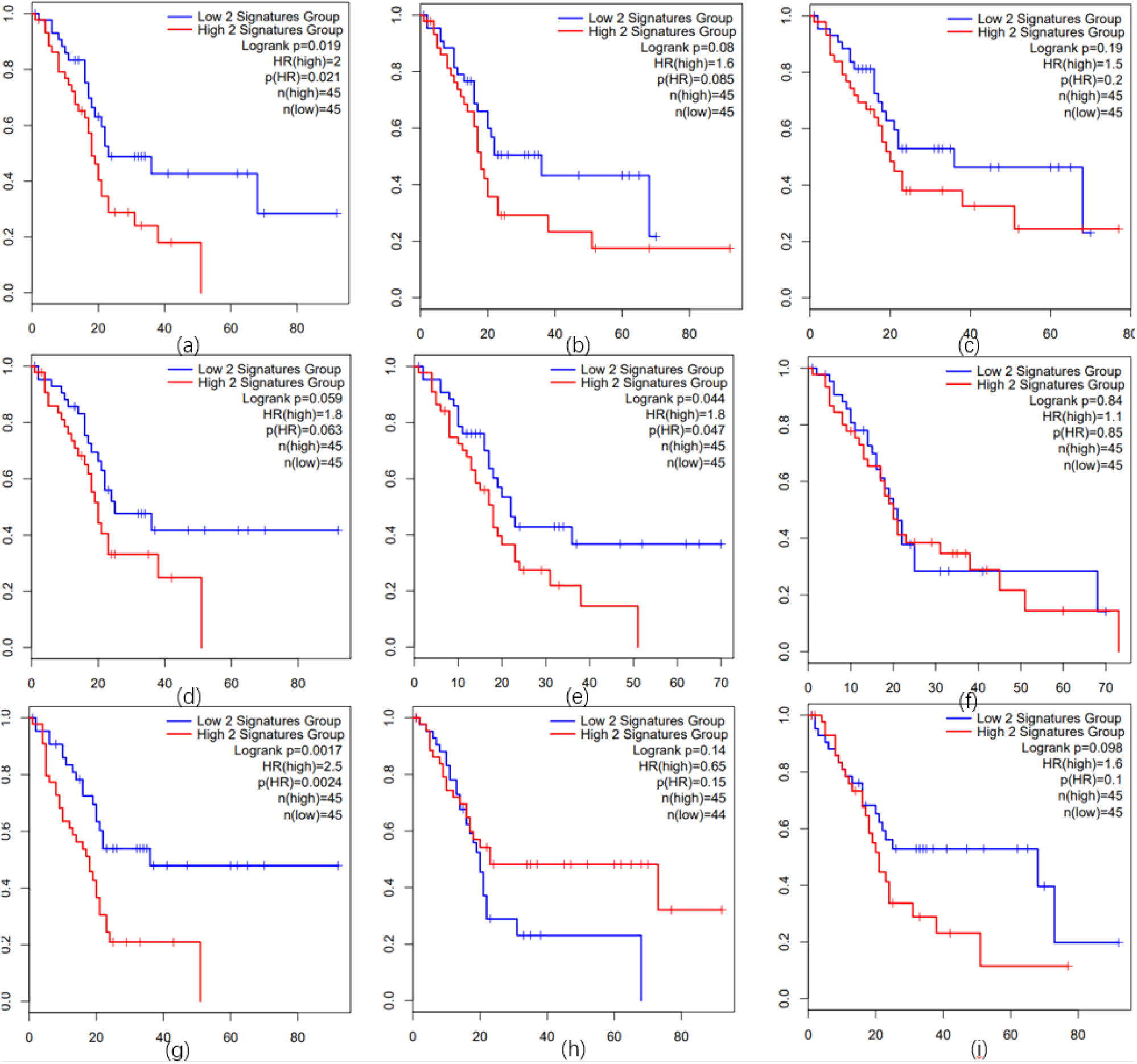
Kaplan-Meier curves for the nine top-ranked target combinations (a)-(i). Kaplan-Meier curves and other survival statistics for (a) < EGLN1, TRFC>, (b) < MAP2K2, TRFC>, (c) < HPSE, TRFC>, (d) < PPIC, TRFC>, (e) < FRK, TRFC>, (f) < EGLN1, COX7C>, (g) < XDH, TRFC>, (h) < MAP2K2, COX7C>, and (i) < FTL, TRFC>. Y-axis indicates survival probability while X-axis indicates months. The blue line in each plot indicates low expression of the two gene groups, and the red line, high expression.

Interestingly, seven of the top nine target combinations include transferrin receptor (TFRC), which encodes a surface receptor responsible for cellular iron intake. High expression of TFRC in PDAC and its strong association with PDAC growth and survival have been reported [36]. Recent studies suggest several key pathways of ferroptosis induction, including mitogen-activated protein kinases (MAPK) and reactive oxygen species (Ros) pathways [37]. Hence, targeting upstream genes (e.g., MAP2K2, EGLN2) along with downstream genes (e.g., TFRC, FTL) might lead to a synergistic effect.

### Performance of DSCNi in predicting drug synergy in cancer cell lines

DSCNi predicts target combinations for individual patients using gene-expression and - essentiality profiles. In this study, we assessed whether DSCNi predicted any association between target- and drug-combination synergy at each individual cell-line level. DrugComb [30] is a comprehensive database that incorporates information regarding the synergy of drug combinations from numerous well-known projects, such as the National Cancer Institute (NCI)-60 [38] for Human Tumor Cell Lines Screen. Because DrugComb includes only one PDAC cell line with five associated combinational drug treatments, we decided to use the cell-line data of triple-negative breast cancer (TNBC). We used 115 TNBC expression profiles from TCGA to generate edge weights in the tissue-function network, 12 TNBC cell lines from the Cancer Cell Line Encyclopedia (CCLE) database [39] to generate edge weights for the cell-line function network, and CRISPR screening data of the TNBC cell line “HS578T” from Project Achilles to generate node weights in the cell-line function network. Among all TNBC cell-lines, HS578T has the largest number (N = 5,226) of drug-combination screening data in the DrugComb database, and our focus on drugs with known targets in DrugBank led to screening data for1,031 drug combinations in the HS578T cell line.) In turn, these drug combinations correspond with 14,066 target combinations in our network model (**Supplementary File 2**).

To measure the association between predicted synthetic lethal pairs and synergistic drug combinations, we constructed a 2 by 2 contingency table (Table 3), in which rows correspond with drug-combination synergy (Y/N), and columns, with target-combination synergy (Y/N). Among synergistic drug combinations, synergy is predicted in 2,594 of their corresponding target combinations with DSCNi, but not in the other 7,097. Neither is synergy predicted in any of the other non-synergistic drug combinations in iDSCN. The *P*-value of the chi-squared test is 0.00001, and the odds ratio is 1,599. This is strong evidence of the greater likelihood that synergistic drug combinations have synergistic target combinations.

**Table 3.** Contingency table between drug- and and target-combination synergy.

|  | Predicted target-combination synergy | Predicted target-combination non-synergy |
| --- | --- | --- |
| Drug-combination synergy | 2,594 | 7,097 |
| Drug-combination non-synergy | 0 | 4,375 |

## Discussion

Our new DSCN method, <u>d</u>ouble target <u>s</u>election guided by <u>C</u>RISPR screening and <u>n</u>etwork, uses both cancer tissue and cell-line models to discover and rank target combinations, and it has several unique features and advantages in comparison with existing methods of selecting combination targets.

For the first time, DSCN uses a subsampling approach that characterizes the knockdown of the first target and models its impact on all the other genes. To demonstrate the validity of this assumption, we studied a set of transcriptome profiles across 22 pancreatic cell lines before and after MAP2K1 and MAP2K2 inhibition. Among 1,301 neighbor genes of MAP2K1 and MAP2K2 in the PPI network, our analysis revealed high correlation of observed log-fold changes in these genes before and after MAP2K1 and MAP2K2 inhibition with log-fold changes calculated from the sub-sampling approach, *R*^2^ = 0.75.

DSCN also differs from all other methods by focusing on the overlapped functional network between cancer tissues and cell lines and further matching the differential gene expression in the tissue to gene essentialities in the cell line. This framework for the selection of target combinations is highly translational and practical. We investigated a number of scoring schemes for calculating impact score, including the most-probable paths, random-walk paths, and diffusion paths, and we studied whether the global network and spectrum clustering-based local network lead to different calculations of impact score. Using tumor samples of pancreatic cancer and cell-line samples and known synthetic lethal data in SynlethDB, we showed statistically significant lower impact scores of target combinations in synthetic lethal gene pairs than other target pairs utilizing a diffusion-path approach on the local network. This analysis clearly demonstrates the validity of our proposed algorithm for calculating the impact scores of target combinations that reflect synthetic lethalilty.

Furthermore, DSCN is broadly defined for every target and target combination, unlike existing network-based target selection algorithms, such as OptiCon[1] or VIPER [2], that are limited by their initial step in the selection of single targets (i.e., master regulators). This advantage of DSCN is demonstrated in the analysis of overlap among the the top-ranked target pairs between DSCN, Opticon, and VIPER and synthetic lethal target pairs reported in the analysis of pancreatic cancer data in SynlethDB. DSCN identified 79 overlapped synthetic lethal target combinations, whereas OptiCon and VIPER showed zero overlap. In addition, three of these top nine predicted synergistic target combinations in pancreatic cancer show statistically significant association with overall survival in patients with pancreatic cancer, and all three contain the TRFC gene, which encodes a surface receptor for cellular iron intake. Hence, the targeting of upstream genes (e.g., MAP2K2, EGLN2) along with downstream genes (e.g., FTL) might lead to a synergistic effect.

Finally, we investigate two relevant but different concepts, drug- and target-combination synergy, hypothesizing the greater likelihood of synergistic than non-synergistic drug combinations to target more synergistic target combinations. Using DSCNi, a model derived from DSCN for the prediction of target combinations for individual patients, we showed the truth of our hypothesis using triple-negative breast-cancer tissue and cell-line data. Based on 1,031 drug combination screening data in HS578T, a TNBC cell line, and its corresponding 14,067 DSCNi-predicted target combination synergy scores, we showed the 1,599-fold higher odds of synergistic than non-synergistic drug combinations to predict synergistic target combinations (*P* = 0.00001).

## Author’s contributions

Enze Liu executed the study. Lei Wang, Xue Wu, Yang Huo and Huanmei Wu provided technical support and valuable comments to the study. Lang Li and Lijun Cheng designed the study.

## Code availability

The python programming was used to implement and train all models. The training and validation datasets used to create each model are available as part of the experimental dataset released as described in materials. The code required to construct the training and validation dataset data and also to analyze the experimental data is provided for download (https://github.com/tzcoolman/DSCN).

